# The role of dietary patterns in the polarization of angiogenic uterine Natural Killer cells during murine pregnancy

**DOI:** 10.1101/2024.07.25.605132

**Authors:** Evila Lopes Salles, Bruno Zavan, Rodolfo Cabral Marcelino, Pablo Shimaoka Chagas, Andrea Mollica do Amarante-Paffaro, Padmashree Chaudhury Woodham, Babak Baban, Valdemar Antonio Paffaro Junior

**Affiliations:** Department of Oral Biology and Diagnostic Sciences, Dental College of Georgia, Augusta University, Augusta, Georgia, USA; Integrative Animal Biology Laboratory, Department for Cell and Developmental Biology, Biomedical Sciences Institute, Federal University of Alfenas, Alfenas, Minas Gerais, Brazil; Department of Food Science and Experimental Nutrition, School of Pharmaceutical Sciences, University of São Paulo, São Paulo, Brazil; Department of Clinical Analyses, Toxicology and Food Sciences, School of Pharmaceutical Sciences, School of Pharmaceutical Sciences of Ribeirão Petro, University of São Paulo, Ribeirão Petro, Brazil; Division of Maternal Fetal Medicine, Department of Obstetrics and Gynecology, Medical College of Georgia, Augusta University, Augusta, Georgia, USA

## Abstract

Uterine Natural Killer (uNK) cells, predominant leukocytes in mouse and human pregnant uteruses, play crucial roles in angiogenesis and pregnancy protection. In mice, DBA lectin-reactive uNK cells expressing Gal-N-Ac sugar exhibit angiogenic functions essential for pregnancy maintenance. This study compares the impact of different nutritional imbalances on mouse pregnancy and the activation of angiogenic DBA+ uNK cells to safeguard against pregnancy complications. High Fat (HF), High Carbohydrate (HC), High Protein (HP), and Food Restriction (FR) diets were administered from gestation day (GD) 1 to GD10 or until parturition. HF and HC diets led to reduced expression of DBA-identified N-acetyl-D-galactosamine, akin to LPS-induced inflammation, and decreased uNK perforin levels. Additionally, HF and HC diets resulted in elevated endometrial cleaved caspase-3 and decreased smooth muscle alpha-actin, causing blood vessel wall thinning without jeopardizing pregnancy term. FR impaired uNK differentiation, manifesting as an “all-or-none” phenomenon with 50% pregnancy failure. Our findings highlight the intricate relationship between nutritional imbalances and mouse pregnancy outcomes. Notably, high-fat diets elicited pronounced responses from DBA+ uNK cells, while high-protein diets had relatively weaker effects. This study underscores the importance of comprehending uNK cell dynamics in maintaining pregnancy homeostasis under diverse dietary conditions, paving the way for elucidating molecular mechanisms governing these interactions. By shedding light on these complex relationships, this research offers valuable insights for improving maternal and fetal health in the context of nutritional interventions during pregnancy.

## Introduction

Changes in nutritional patterns are known to influence metabolism, with overnutrition increasing energy expenditure and undernutrition reducing it [1, 2]. Despite being opposite conditions, both can have detrimental effects on reproductive capacity [3, 4].

Maternal undernutrition presents a significant risks of pregnancy complications and poor fetal development.[5, 6]. As with undernutrition, overnutrition negatively affect fertility [7]. Obesity-related pregnancy complications increase the risk of preterm birth, miscarriage, gestational diabetes, and hypertensive disorders and fetal programming alterations, leading to long-term health issues in offspring [8–10].

Normal pregnancy is characterized as an immunosuppressive state, with a high number of uterine Natural Killer (uNK) cells producing of Interferon-gamma (IFN-γ), essential for pregnancy-induced spiral artery remodeling and placental development [11]. In pregnant mice, uNK cells rapidly increase in number and size and acquire granules until gestational day (GD) 10, forming the transient endometrial structure known as the mesometrial lymphoid aggregate of pregnancy (MLAp) [12].

Subsequently, uNK cell numbers gradually decline until term, accompanied by nuclear fragmentation [13, 14]. During mid-pregnancy, the majority of mouse uNK cells express a surface N-acetyl-D-galactosamine (GalNac) sugar, selectively marked by Dolichos biflorus agglutinin (DBA) lectin histochemistry, allowing the characterization of four maturation-related subtypes of DBA^+^uNK cells [15].

DBA^+^uNK cells predominantly express transcripts for angiogenic factors [16] and, although poorly cytotoxic these cells containing granules encasing perforin and granzymes [17, 18]. About 95% of the DBA^+^uNK cells can be found in the uterus, precisely within the area of pregnancy-associated neovascularization [19, 20].

This study aimed to prospectively investigate the impact of potential stressful and/or immune-inflammatory unbalanced diets on a mouse pregnancy experimental model, focusing primarily on the angiogenic DBA^+^uNK cell analyses. We hypothesized these nutritional alterations could impact these cells found in mouse and human uterus during pregnancy.

## Materials and methods

### Animals

Female SWISS Webster mice (8-10 weeks old) were mated with SWISS males, and the presence of a copulation plug was considered as GD1. The mice were housed in the Central Animal Facility of the Federal University of Alfenas (Unifal-MG, Brazil) under controlled conditions of light (12:12h light-dark cycle) and temperature (23±1oC), with ad libitum access to food and water, except those subjected to food restriction (FR). A total of 90 mice were included in the analysis, and all animal procedures were following the U.K. Animals (Scientific Procedures) and approved by the local ethics committee (Protocol number: 448/2012).

### Diets

The diets were developed by the In vivo and in vitro Nutritional and Toxicological Analysis Laboratory (Lantin) at UNIFAL-MG. At the GD1, pregnant females were assigned to one of the following groups: Control (CD), High Protein (HP), High Fat (HF), High Carbohydrate (HC), or a Food Restriction (FR) diet, where animals received 4g of feed/day (Figure 1). The detailed feed composition of the diets is presented in supplementary data (Supplementary figure 1). From GD1 to GD10, all females were weighed, and the food intake was monitored.

**Figure 1.**
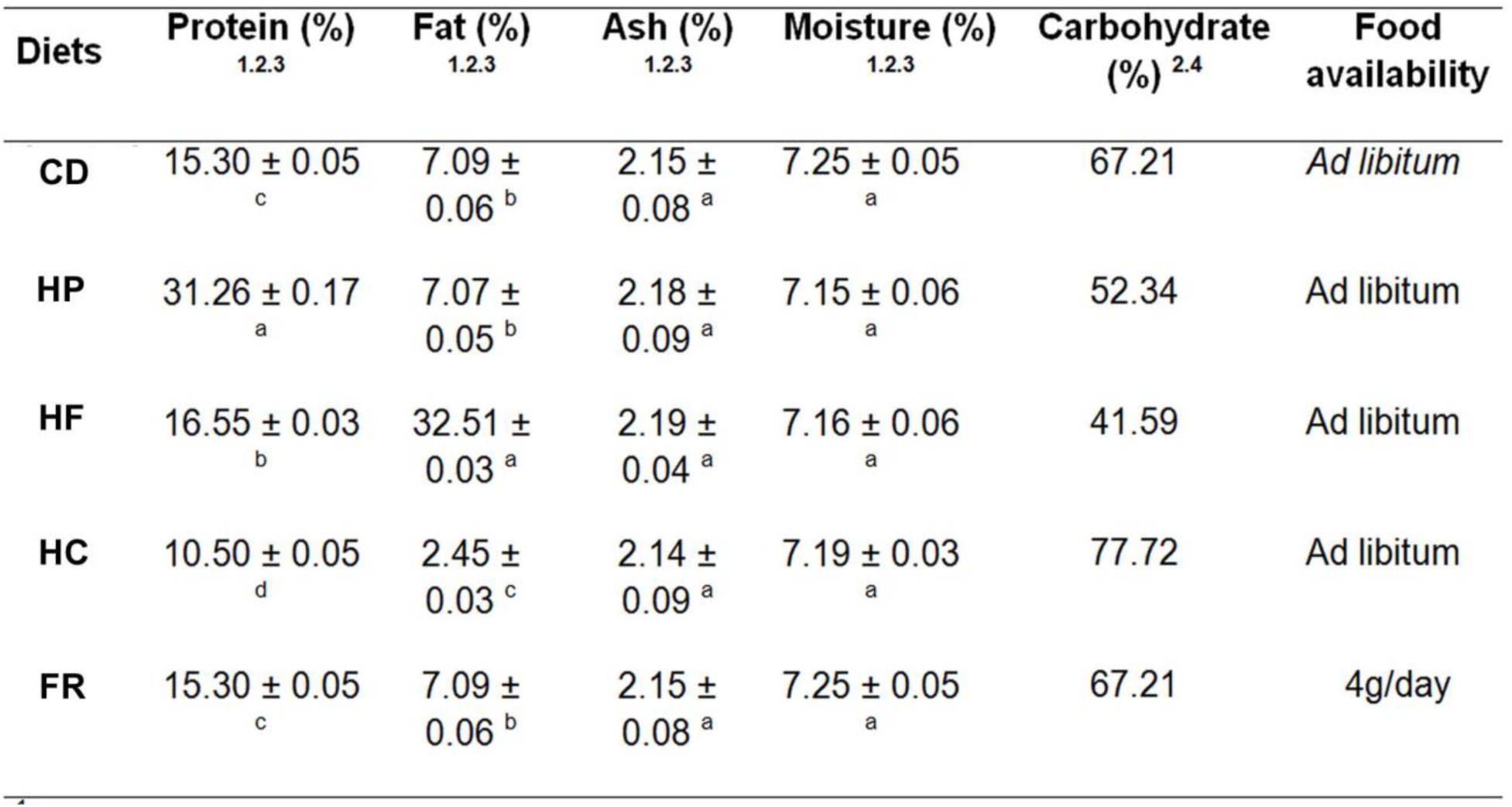
Centesimal composition of the experimental diets. 1. Values correspond to means (± SD) of three determinations; 2. Values expressed in dry basis; 3. Values not sharing similar letter in the same column are different (p < 0.05) in Tukey test; 4. Calculated by difference = 100 – (protein + total fat + ash + moisture). Control Diet (CD). High Proteic Diet (FR). High Fat Diet (HF). High carbohydrate diet (HC) and Food restriction diet (FR).

### Pregnancy viability and litter analysis

On GD10, sixty-five females were anesthetized with 2% inhaled Isoflurane (BoChimico, Itatiaia, RJ, Brazil) and perfused with 4% paraformaldehyde (Sigma-Aldrich, St. Louis, MO, USA) in PBS (50mM). The developing implantation sites and/or reabsorbed implantation sites were macroscopically analyzed to assess pregnancy viability (CD: n=10 mice; HP: n=10; HF: n= 10; HC: n=14 and FR: n=21). Five randomly selected pregnant females from each group had their implantation sites dissected and subjected to Hematoxylin and Eosin (H&E) staining, DBA Lectin, α-actin, perforin and cleaved caspase-3 labeling. Histological sections stained with H&E were analyzed under light microscopy to verify the incidence of implantation sites under resorption and possible dietary-induced morphological alteration. Litter size and pup weight analyses were performed on 25 animals fed the diets until full term pregnancy (5 animals/group).

### DBA lectin histochemistry

Histological sections from five mice per group were deparaffinized, hydrated, and subjected to DBA lectin histochemistry as described by Paffaro et al., 2003. The sections were then examined under light microscopy (Nikon Eclipse 80i, Tokyo, Japan).

### Stereological and Morphometric Study

Three histological mid-sagittal sections (7µm) labbeled with DBA lectin from three implantation sites of five animals from each experimental group were used for stereological analysis. The density profiles (QA) of the four morphological subtypes of DBA^+^uNK cells were determined based on cells size, chromatin condensation, and N-acetyl-galactosamine expression on their cell surface and in granules. The diameters of the four uNK cells subtypes were measured (50 cells/subtype/group) and counted by two experienced observers in three regions of the implantation site on GD10 (Figure 4). In these regions, three test areas (TA) of 4.104µm2 were used to quantify the subtypes, with TA defined as a quadratic test system with two exclusion lines, including only cells with visible nucleus.

Arteriole wall and DB+MLAp morphometry was assessed in implantation sites (n= 5 animals/group) under light microscopy (Níkon Eclipse 80i, Tokyo, Japan) using image analysis software (NIS-Elements/Nikon/Japan). The ratio of the Total Area to Luminal Area was determined from 300 arterioles (75 arterioles/group). The total area of DB+MLAp was also measured in the same histological slides.

### Perforin, α-actin and cleaved caspase-3 immunohistochemistry

Histological sections from implantation sites (n=5 animals/group) were deparaffinized, hydrated, and subjected to immunohistochemistry. For Perforin and α-actin analyses, sections were submitted to 1% hydrogen peroxide (Sigma-Aldrich, St. Louis, MO, USA) for 30 min. After washing with 50mM PBS, sections were incubated with 1% bovine serum albumin (BSA) (Sigma, St Louis, MO, USA) in PBS for 30 min, followed by overnight incubation at 4°C with rabbit-primary antibodies anti-mouse Perforin (1:50) (PA5-17431, Thermo Scientific. USA) or anti-mouse α-actin (1:100) (A2103, Sigma-Aldrich. MO.USA) over-night at 4°C. Sections were subsequently incubated with biotinylated anti-rabbit secondary antibody (1:500) (B8895 Sigma-Aldrich. MO.USA) for 60 min at room temperature. Subsequently, sections were washed in PBS and incubated with RTU Horseradish Peroxidase Streptavidin (SA-5704, Vector Laboratories, Burlingame, CA) for 1 hour at room temperature and 3,3-diaminobenzidine (Sigma, St. Louis, MO, USA) in 50mM TBS containing 0.1% hydrogen peroxide. Sections were counterstained with Harris’s hematoxylin, mounted with Entellan (Merck, Darmstadt, Germany) and observed under light microscopy (Nikon Eclipse 80i, Tokyo, Japan). For cleaved caspase-3 immunostaining, the deparaffinized and hydrated sections were washed with 0.5M PBS, pH 7.4, followed by blocking of unspecific binding sites with 1% PBS/BSA (Bovine serum albumin-SIGMA) for 30 min. Sections were then incubated with anti-cleaved caspase-3 antibody (1:50, MI0035, Rhea Biotech, BRA) overnight at 4°C. Subsequently, sections were incubated for 120min with anti-rabbit FITC secondary antibody (1:250, F0382, Sigma-Aldrich. MO.USA) and 4’,6-diamidino-2-phenylindole (DAPI, 1:1000, D9542, Sigma-Aldrich. MO.USA) for 5 min.

### Staining quantification by Pixel density

From histological sections submitted to DBA lectin histochemistry, anti-Perforin, anti-α-actin and anti-cleaved caspase-3 investigation, at least five images/captured at 100x magnification were analyzed for pixel density using GNU Image Manipulation Program (GIMP 2.8.10 software) as previously described. During pixel density analyses, careful examination was done to avoid technical artifacts, such as over-development and precipitation in all sections.

### Statistical analysis

After analyzing the data for column homoscedasticity, we performed a one-way ANOVA test, followed by post-test analysis. Weight differences were analyzed by Student’s t test. Statistical significance was set atp≤0.05.

## Results

### Food Intake and weight analyses

The HC-diet mice exhibited increased food intake compared to CD-diet mice on GD2 (p<0.001). However, no difference in food intake was observed between the groups on GD3 (p>0.05). The FR group received 4g of the control diet daily, and the total consumption of the 4g feed by all mice from this group was observed every morning (Figure 2A).

**Figure 2.**
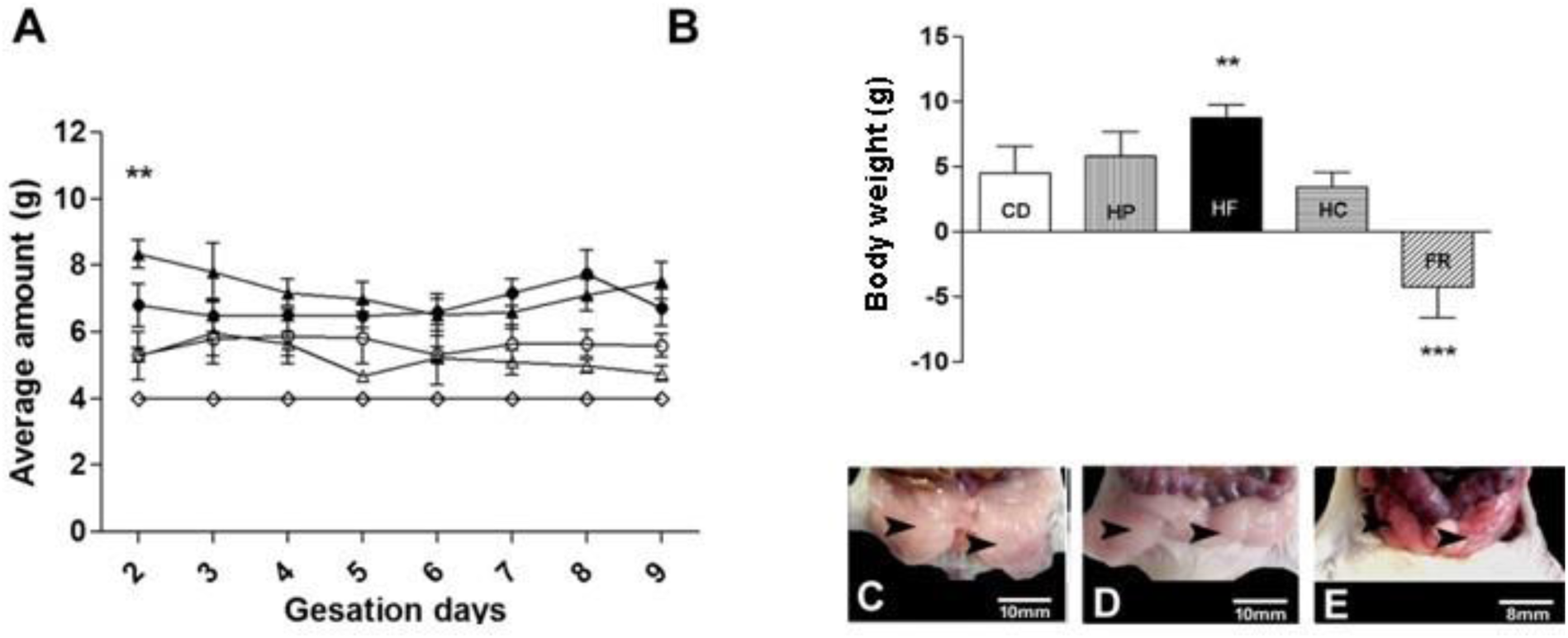
Food intake analyses (A). High carbohydrate diet [▴] High Protein diet [●]. High HF diet [Δ], Calorie diet [○] and Food Restriction [◊]. Body weight gain analyses (B). Control Diet (CD). High protein diet (HP). High Fat diet (HF). High Carbohydrate diet (HC). Food restriction diet (FR). p≤0.01 (**), p≤0.001(***). Macroscopic images of perigonadal adipose tissue (arrow heads) from CD-fed mice (C), HF-fed mice (D) and FR-fed mice (E).

Evaluation of weight gain from GD1 to GD10 showed that mice fed the HF diet exhibited a significant weight gain (7.31g, p<0.001) compared to control diet (Figure 2B). HF diet-fed mice also exhibited a greater amount of visceral adipose tissue (VAT) (Figure 2C, Figure 2D) while FR diet-fed mice exhibited less VAT (Figure 2E) and weight loss (-4.25g, p<0,0001). However, no differences in weight gain were found in mice fed the HP (5.82g) and HC (2.62g) diets (p>0.05) compared to the control (4.51g).

### Implantation Site Analysis

The IS from all animals analyzed were macroscopically evaluated and counted when pregnancy was confirmed during laparotomy on GD10. There were no differences (p>0.05) among pregnant mice fed on HP (13.27 IS/mouse), HF (13.92 IS/mouse), HC (14.54 IS/mouse) and FR (14.30 IS/mouse) compared with CD-diet mice (14.15 IS/mouse) (Figure 3A). However, four animals from HC group and 11 animals from FR group did not exhibit IS after laparotomy and so they were considered not pregnant (Figure 3B), while animals from CD, HP and HF groups exhibited 100% pregnancy rates.

**Figure 3.**
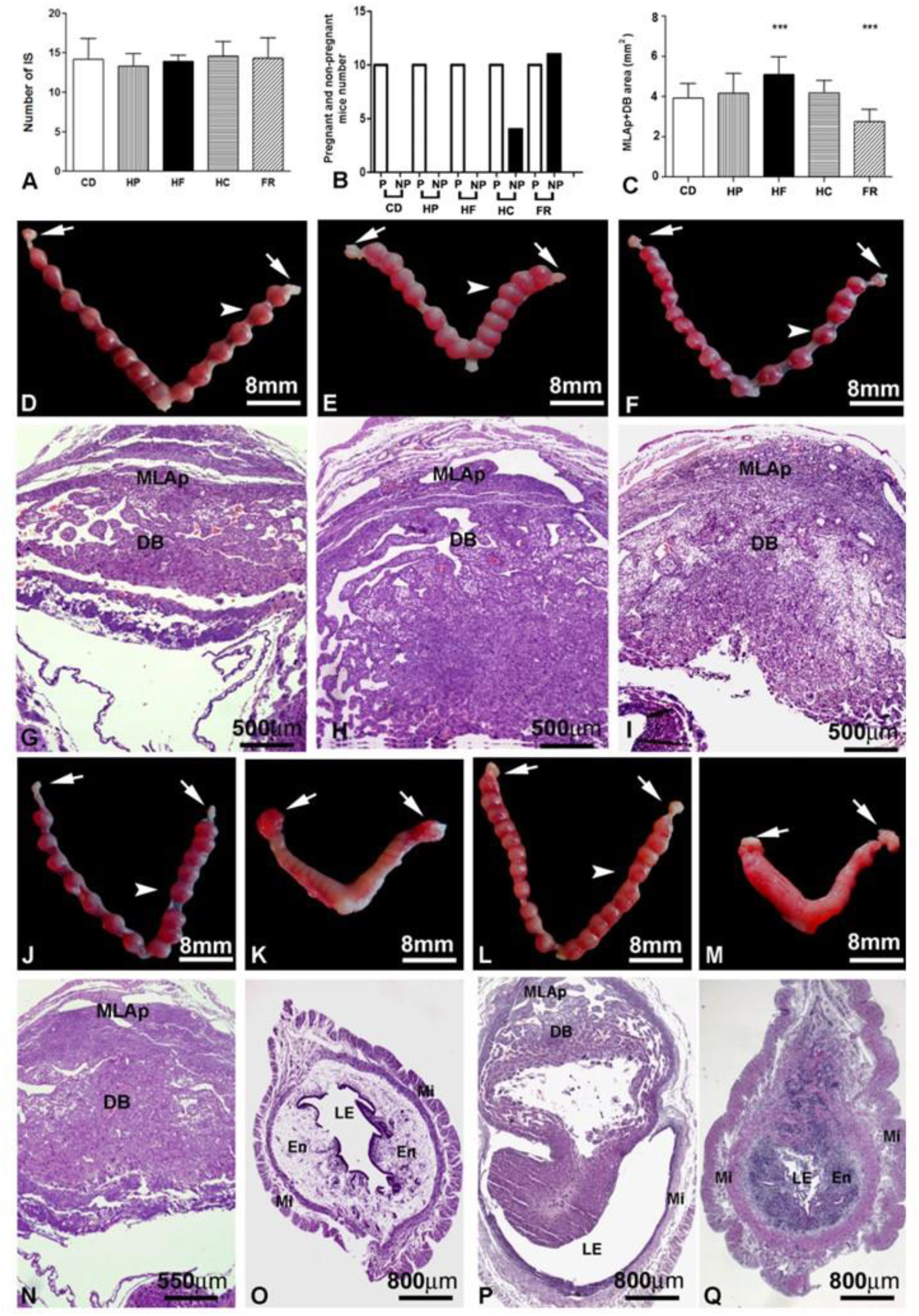
Study of gestational viability on the GD10 showing number of implantation sites (A), pregnancy rate (B), Mesometrial lymphoid aggregate of pregnancy (MLAp) plus Decidua Basalis (DB) area (C). p≤0.001 (***). Macroscopic images show examples of implantations sites from CD (D), HP (E), and HF-fed mice (F). Photomicrographs of histological sections from CD (G), HP (H) and HF-fed mice (I). Macroscopic images show examples of implantation sites from HC (J and K), Food restriction mice (L-M). Photomicrographs of histological sections from HC (N and O), Food restriction (P and Q). Note examples of non-pregnant uterus only from HC (K and O) and from FR mouse (M and Q). In P, note the resorption aspect of the embryo implantation site. Ovaries (Arrow). Implantation sites (arrowhead). Luminal epithelium (LE), Endometrium (EN). Myometrium (My).

Macroscopic analyses of mice that were fed on CD, HP, HF, HC and FR diets showed uterine horns containing IS with regular morphology and without resorption or hemorrhagic sites (Figures 3D, 3E, 3F, 3J, 3L). Microscopic analyses showed regular IS histoarchitecture from CD, HP, HF, and HC-diet fed mice. In those IS, it was possible to identify the large MLAp and Decidua Basalis (Figure 3G, 3H, 3I, 3N). About 2 IS/mice resembling resorptions were observed in histological sections of pregnant FR diet-fed mice. In these animals, a disorganized uterine histoarchitecture, a large lumen, and hemorrhagic sites containing cells with pyknotic nuclei were identified (Figure 3P). Microscopic evaluation of non-pregnant HC and FR diet-fed mice uterus showed the regular virgin uterine morphology (Figure 3O, 3Q) consistent with macroscopic analyses (Figure 3K, 3M).

### DBA lectin histochemical analysis

DBA lectin histochemical analysis showed GalNac expression (DBA+ reaction) in all IS analyzed. DBA^+^ reactions were localized on uNK cell that were distributed at the three regions of the IS (Figure 4A, 4B, 4C, 4D, 4E). In the CDS group, DBA+ reactions were localized on the uNK plasma surface and granules which allowed the identification of four morphological DBA^+^uNK subtypes (Figure 4F) [15, 21]. We observed in IS from HF and HC diet-fed mice several uNK cells that had low expression of GalNac sugar (DBA^low^uNK). These uNK cell subtype exhibited low DBA lectin reactivity on their surface and/or within their granules (Figure 4G). The lower DBA reactivity was confirmed also by semi-quantitative analysis (Figure 4H) which showed GalNac expression was significantly weak (p≤0.05) in the IS from mice that were fed on the HF (Figure 4C) and HC (Figure 4D) diets in comparisonwith the control.

**Figure 4.**
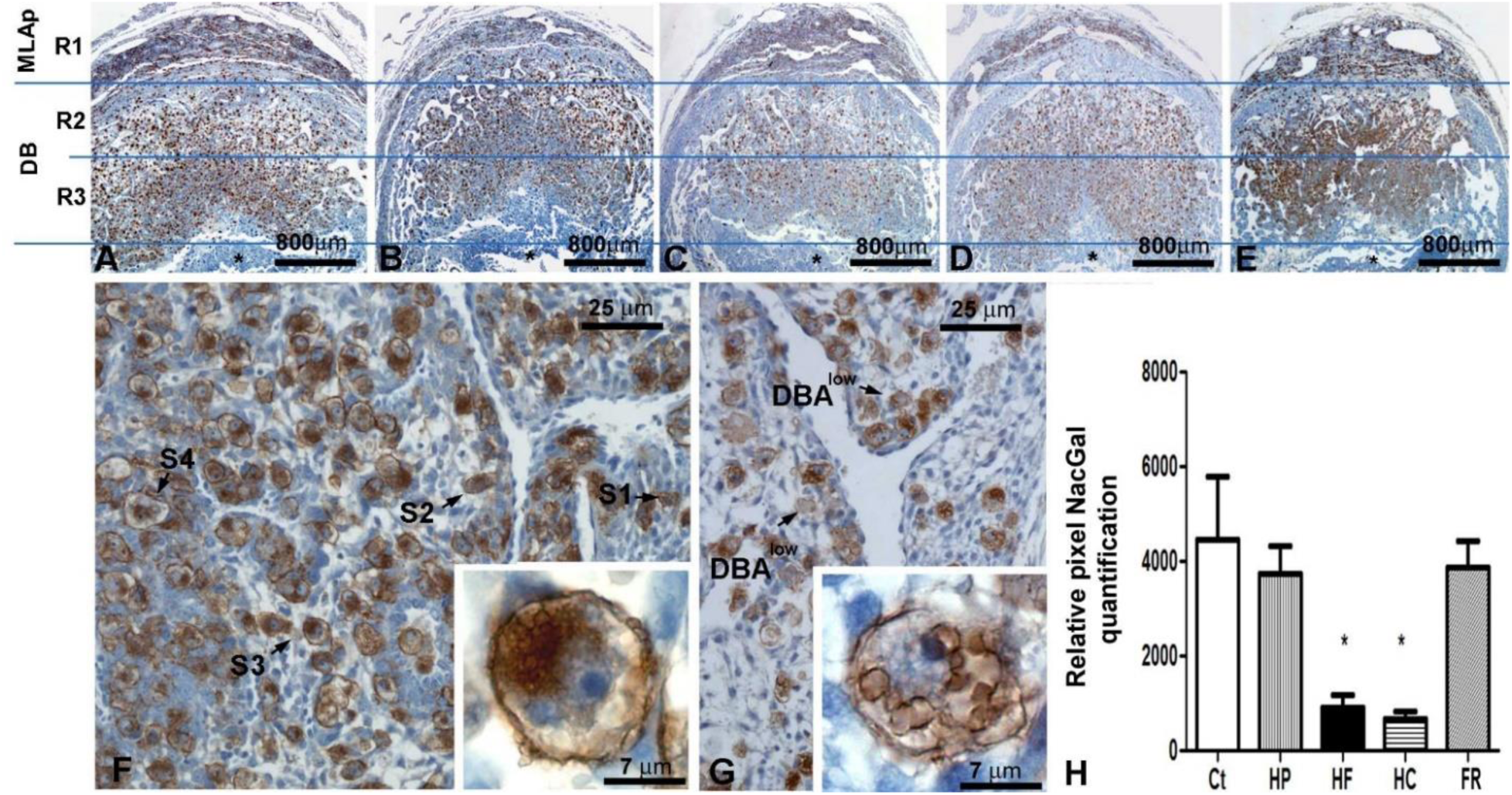
Photomicrographs of implantation sites (IS) from pregnant mouse uterus on GD10. Note the regions that were used to uNK cells quantification (R1, R2 and R3) in the panoramic pictures from these IS (A-E). Decidua Basalis (DB). Mesometrial lymphoid aggregate of pregnancy (MLAp). Observe the strong DBA reaction in the mice fed with CD (A), HP (B) and FR (E) diets compare to the weak DBA reaction observed in mouse fed HF (C) and HC (D) diets. Detail of the DBA lectin reaction pattern found in IS from CD fed mouse (F). Note subtype 1 (S1), Subtype 2 (S2), Subtype 3, (S3) and Subtype 4 (S4) uNK cells. Insert in F shows the same S3 uNK cell as a large and high granulated cell exhibiting euchromatin predominant in the nucleus and nucleoli. Detail of the DBA lectin reaction pattern found in IS from HF fed mouse (G). Note the weak reaction in several uNK cells (DBA^low^). Insert in G shows DBA^low^ uNK cell subtype exhibiting irregular DBA lectin reaction in the surface and several large empty-like granules, nucleus with euchromatin predominant and nucleoli. Relative pixel NacGal-DBA lectin detected quantification (H). p≤0.05 (*).

### Morphometric and Stereological analysis of DBA^+^uNK

To address the effect of different diets on DBA^+^uNK from pregnant mice, we carefully analyzed, measured, and quantified the four subtypes of DBA^+^uNK cells and the DBA^low^uNK subtype at three regions from histological sections of implantation sites.

The number of the subtype I of DBA^+^uNK cells had not changed in any of the three regions of all the five experimental groups. This uNK cell subtype was abundant at region 1 of IS from mice fed on HP, HF, HC, and FR diets similar to control diet-fed mice (Figure 5A, 5B, 5C). However, they were bigger in HC and HP diet-fed mice compared with control (Figure 5D).

**Figure 5.**
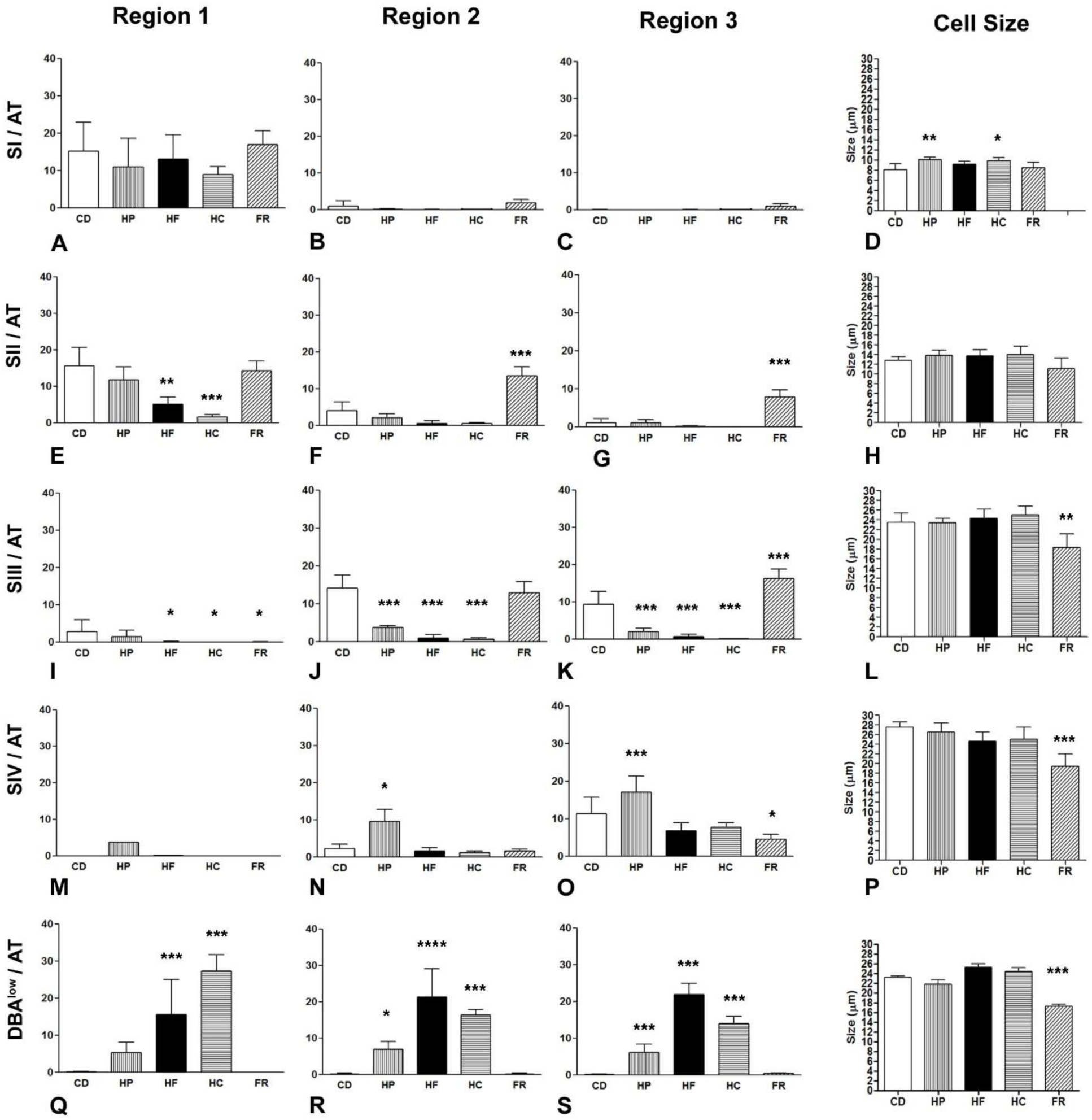
Stereological and morphometric uNK cell analyses. Quantification and cell size measurement of four DBA+ uNK cell subtypes (S1-S4) and the DBA^low^ uNK cell (DBA^low^) in each of the three regions of the GD10 embryo implantation site from each experimental group. Control diet fed mice (CD), High Protein diet fed mice (HP) High Fat diet fed mice (HF), High Carbohydrate diet fed mice (HC), Food restriction diet fed mice (FR). p≤0.05 (*), p≤0.01 (**), p≤0.001(***), p≤0.0001(****).

The subtype II of DBA^+^uNK cells decreased in number at IS regions 1 in HF and HC diet-fed mice. However, in FR diet-fed mice at IS region 2 and 3, the number of subtype II was higher (Figure 5F and 5G).

The number of subtype III DBA^+^uNK cells were different in all experimental groups compared to control. In mice fed with HF, HC, and FR the number of these cells was reduced at IS regions 1, 2 and 3. While HP group showed reduced number of these cells only in regions 2 and 3 (Figure 5E, 5F and 5G). At region 3 from FR fed-diet mice (Figures 5I, 5J, 5K), the subtype III of DBA^+^uNK cells was higher than the same region of CD group. Subtype III cells presented smaller diameter in FR diet-fed mice compared to control group at the same region (Figure 5L).

Subtype IV statistically increased in IS from mice HP diet-fed in all the implantations sites regions (1, 2 and 3) compared with control group. In FR mice, however, the number of Subtype IV was lower at region 3 (Figures 5M, 5N, 5O). Also, these cells were smaller in FR diet-fed mice than in control group (Figure 5P).

The DBA^low^ uNK subtype was found through all regions of IS from HP, HF, and HC mice (Figures 5Q, 5R, 5S), but were rare in FR (Figure 5T) and control mice. Also, the DBA^low^ uNK found in FR were smaller than the ones found in the other groups.

### Immunohistochemistry to perforin and cleaved caspase-3

As expected, a strong well-localized and specific perforin reaction was observed in cytoplasmic granules of DBA^+^uNK cells at region 1 (Figure 6A), region 2 (Figure 6B) and region 3 (Figure 6C) of IS from control mice.

**Figure 6.**
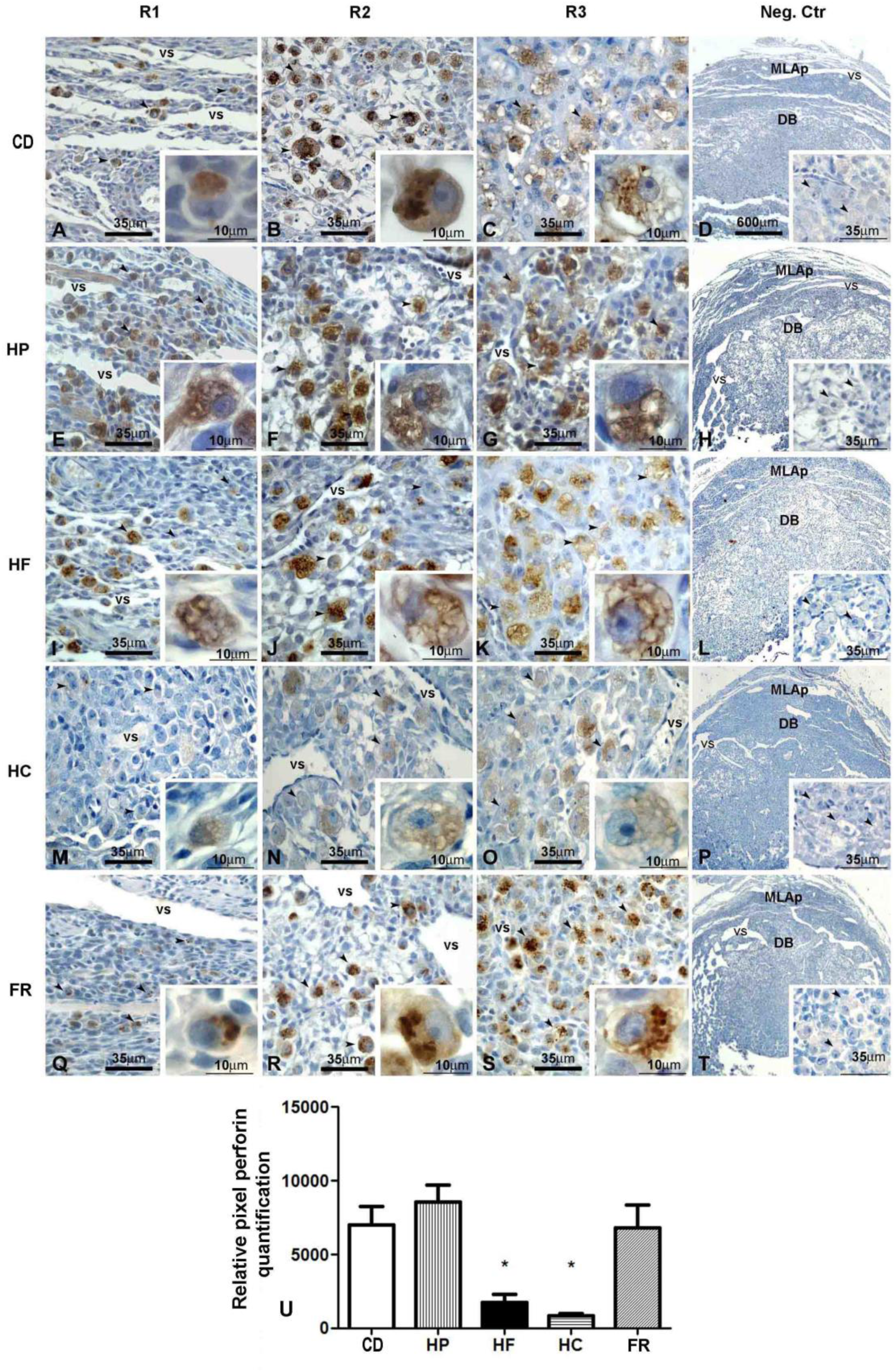
Photomicrographs of Perforin immunocytochemistry analyses in each of the three regions (R1, R2 and R3) of the GD10 embryo implantation site from each experimental group (A-T). Control diet fed mice (CD), High Protein diet fed mice (HP) High Fat diet fed mice (HF), High carbohydrate diet fed mice (HC), Food restriction diet fed mice (FR). Observe the weak perforin reaction in the images from HF and HC mice compared to the CD mice. Negative control for perforin (Neg.Ctr). Inserts show high magnification of uNK cells and their reactivity to the perforin antibody in all 3 regions analyzed. Mesometrial lymphoid aggregate of the pregnancy (MLAp). Decidua Basalis (DB). Relative pixel perforin quantification (U). p≤0.05 (*).

In females fed with HP diet, despite the uNK cells present reactive to anti-perforin antibody, the reaction was apparently weak and was not well-localized in their granules (Figure 6E, 6F, and 6G). In IS of HF diet-fed mice, the granules from uNK cells resemble empty granules and the perforin positive reaction were observed delineating the granules similar to the DBA^low^uNK cell subtype observed in our histochemical studies (Figures 6I, 6J and 6K). The same happened at IS from HC fed-diet mice (Figures 6M, 6N, and 6O). However, IS of FR females showed strong anti-perforin staining in the granules of uNK cells in a similar well localized manner observed in the control (Figure 6Q, 6R, 6S). Quantitative densitometry showed a significantly lower anti-perforin reaction in IS on mice fed on the HF (p≤0,05) and HC (p≤0,001) diets (Figure 6U) compared with the control.

The analysis of the fluorescent staining for cleaved caspase-3 showed a weak reaction in IS from control (Figures 7A, 7F), HP (Figures 7B, 7G) and FR-fed mice (Figures 7C, 7H), while in IS from HF (Figure 7D, 7I) and HC (Figures 7E, 7J) fed mice the reaction was strong. Unfortunately, the exact cell types reacting with the anti-caspase 3 antibody were not identified in our study. The quantitative densitometry analyses of the immunoreaction confirm data obtained under microscopy (Figure 7P).

**Figure 7.**
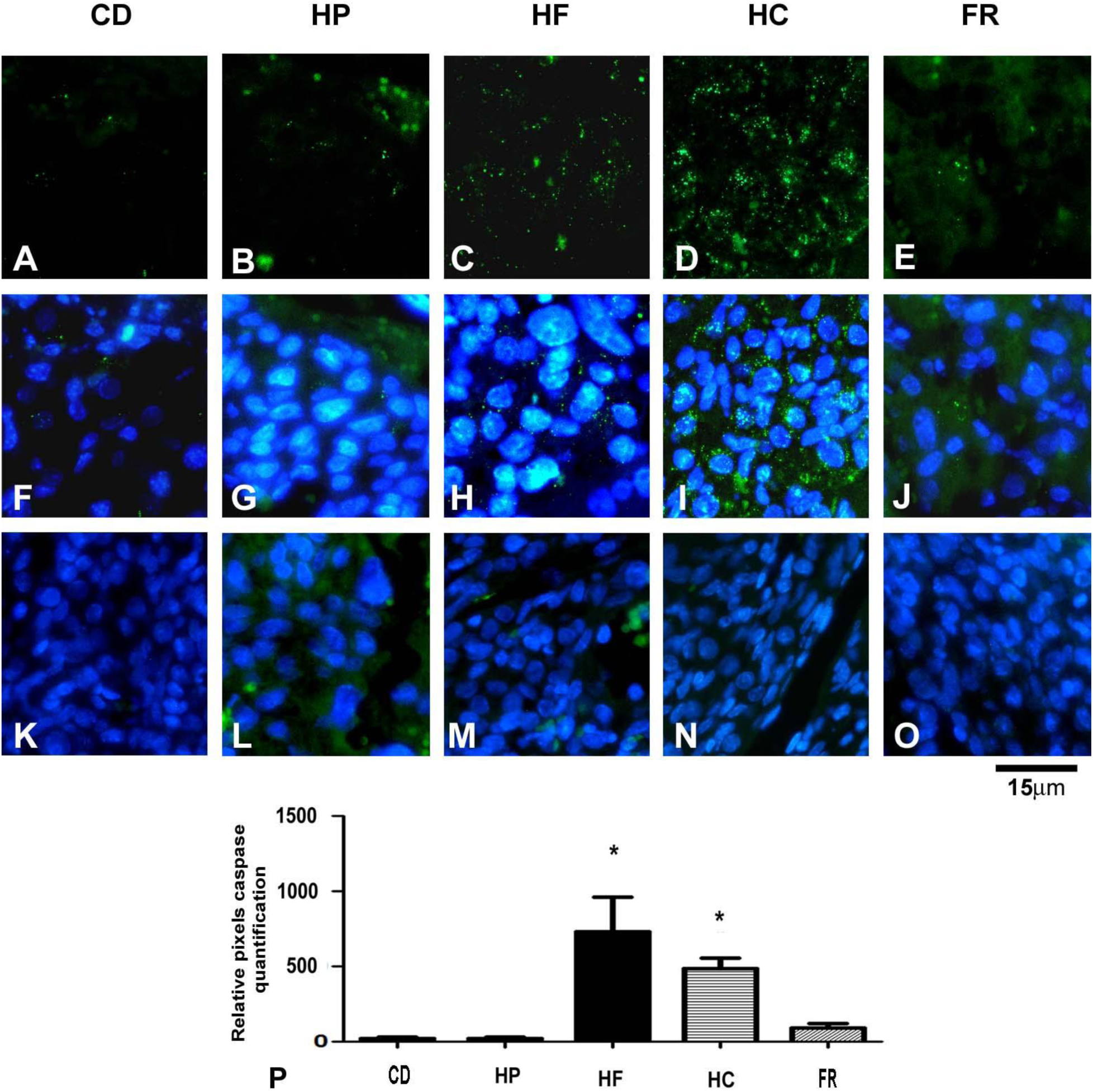
Immunofluorescent photomicrographs of the GD10 embryo implantation site from each experimental group submitted to the 3 cleaved caspase. 3-cleaved caspase (Green) in control diet fed mice (B), High protein (C), High Fat (D), High carbohydrate (E) and Food restriction (F). 3 cleaved caspase (Green) and DAP (Blue) in control diet fed mice (G), High protein (H), High Fat (I), High carbohydrate (J) and Food restriction (K). Examples of 3-cleaved caspase negative control in each treatment respectively (K-O). Relative 3 -cleaved caspase quantification (A) p≤0.05 (*).

### Alpha-actin immunohistochemistry and Morphometric analysis of uterine arteries

In control mice we, observed strong α-actin staining inside uNK cells, which were morphologically normal (Figure 8A). In addition, a strong alpha-actin positive reaction was found in the smooth muscle of blood vessels from the endometrium (Figure 8B) and in smooth muscle cells in the myometrium (Figure 8C). The same was observed in HP (Figure 8E, 8F and 8G) and FR fed mice (Figures 8Q-8T). In HF (Figure 8I) and HC (Figure 8M) fed-diet mice sections, a weak positive reaction was observed in cells resembling uNK and at blood vessels (Figure 8J and 8N), but a strong myometrial reaction could still be observed in these groups (Figure 8K and 8O).

**Figure 8.**
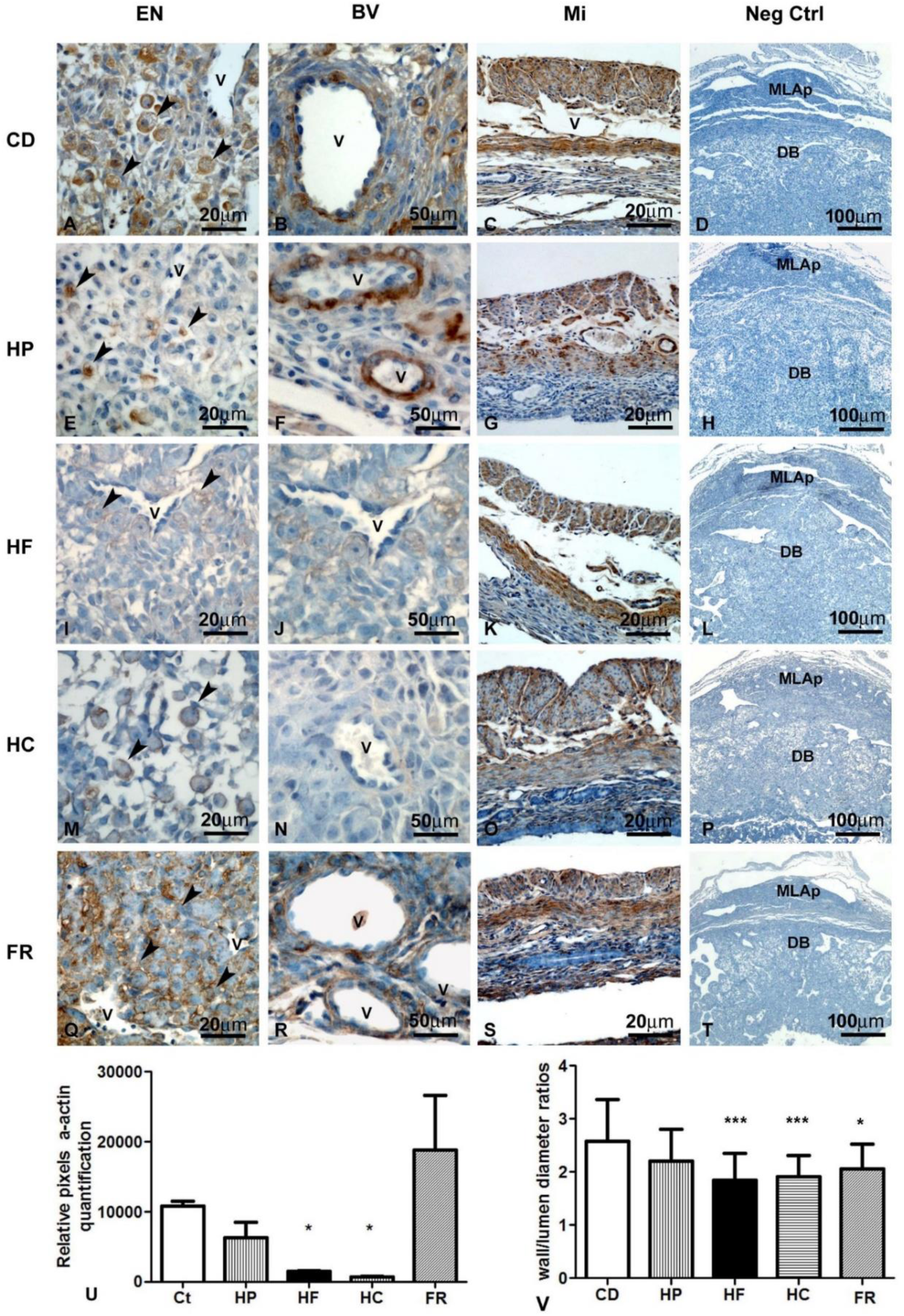
Photomicrographs showing examples of implantation sites alpha-actin immunohistochemistry analyses. Endometrium (A), detailed blood vessels (B), myometrium (C) and Negative control reaction (D) from mice ed control diet. Endometrium (E), detailed blood vessels (F), myometrium (G) and Negative control reaction (H) from mice fed High Protein diet. Endometrium (I), detailed blood vessels (J), myometrium (K) and Negative control reaction (L) from mice fed High Fat diet. Endometrium (M), detailed blood vessels (N), myometrium (O) and Negative control reaction (P) from mice fed High Carbohydrate diet. Endometrium (Q), detailed blood vessels (R) myometrium (S) and Negative control reaction (T) from mice fed Food restriction diet. Relative blood vessels alpha actin quantification (U). Morphometric blood vessels analyses (V) p≤0.05 (*), p≤0.001 (***).

The weak blood vessels’ reaction from HF and HC were confirmed through relative pixel quantification (Figure 8U). The analyses of arteriole wall thickness showed no difference (p≥0.05) between control (r=2,58) and HP fed-diet mice (r=2.2). Regardless the HF (r=1.8; p≤0.0001) HC (r=1.8; p≤0.0001) and FR (r=2 and p≤0.0001) fed mice showed thinner arteriole walls in comparison with control mice (Figure 8V).

### Litter size and weight analysis

The present study showed a litter size reduction from FR group (4.2 pups), while HP (10.3pups), HF, (12.6 pups) and HC (12.3) fed-diets mice were similar to control (14 pups) (Figure 9A). Pup weights were also lower in FR (2.8g) and HP (3g) fed-diets mice compared with control (3.3g) (Figure 9B). The animals that carried gestation to term, the birth of offspring occurred within 20 days after copulation.

**Figure 9.**
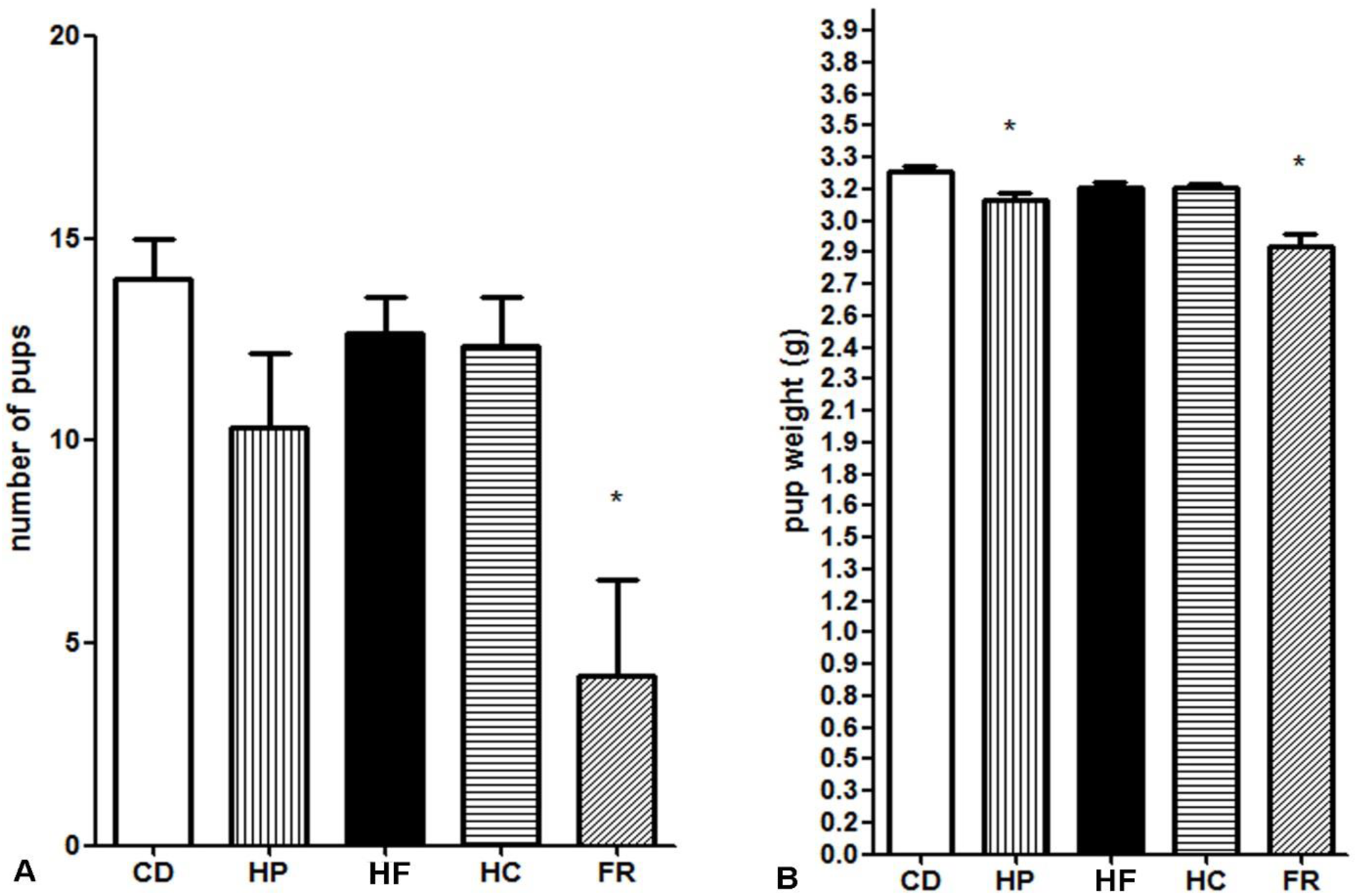
Pregnancy to term study in each experimental group: control diet fed mice (CD), High protein (HP), High Fat (HF), High carbohydrate (HC) and Food restriction (FR). Number of pup quantification (A). Pup weight (B).

## Discussion

Uterine Natural Killer cells are essential for proper uterine spiral artery remodeling, fetal and placental development, and the overall success of pregnancy. Our study elucidated how high intake of different macronutrients and food restriction can impact angiogenic uNK.

The pregnancy viability analysis showed that the FR and HC diets caused total pregnancy failure in 50% and 25% of the mice respectively. The phenomenon of pregnancy failure under food restriction has been previously documented in mice [22] and rats [23, 24]. Pascalon and Bertrand (1987) [24] named t effect of FR on pregnancy viability as an “all-or-none” phenomenon, where (i) females either fail to maintain pregnancies, or (ii) carry them to term with full-size litters; as observed in our study.

Although the phenomenon has been described, its implications for IS haven not been deeply explored. There are no data in the literature regarding the effects of FR on the angiogenic uNK cells activity. However, our analysis showed that FR led impaired decidua/MLAp and uNK cells development, along with an increase of subtypes II and III cells, a decrease in subtype IV cells and smaller cells. The restriction of nutrients delayed the angiogenic uNK cells development and contributed to the decreased wall/lumen diameter of decidua vessels. These results evidently impacted the pregnancy viability in FR group, as the density and activation of uNK cells are associated with pregnancy complications such as fetal growth restriction and recurrent spontaneous abortion [25], both outcomes observed in our study.

The HC group had 25% pregnancy failure rate, and the pregnant females t showed an increase in number of DBA^low^uNK cells and fewer subtypes II and III uNK cells. DBA^low^uNK cells were also present in the HF diet group. The similarity between these results is due to the similar effects induced by high fat and carbohydrate intake, both of which have been shown to trigger inflammation [26]. Interesting, DBA^low^uNK cells were first described in 2015 after the administration of LPS in pregnant mice [27].

It has been reported that LPS injection in pregnant mice induces down-regulation of alpha-actin and perforin loss in uNK cells [27], similar to what was observed in IS from HC and HF groups. The molecular basis described in the literature support the hypothesis that inflammation is the underlying cause of the outcomes observed in HC and HF females. Both LPS and HC or HF diets can trigger inflammation through the activation of common pathways such as NF-kB signaling [28, 29], the release of pro-inflammatory cytokines such as tumor necrosis factor-alpha (TNF-α), [30, 31], and the promotion of tissue-specific inflammation [32–34].

Another point of consideration is the effect of HC and HF diets on leptin levels. Levels of leptin increase significantly during pregnancy [35] and play a crucial role in regulating conceptus growth and development, fetal/placental angiogenesis, and other events [36]. High intake of fat and carbohydrates also induces higher leptin concentrations [37]. Leptin action contributes to chronic inflammation in obesity [37]. Maternal obesity, which can lead to elevated leptin levels, has been linked to altered uNK activity through a functional imbalance of killer immunoglobulin-like receptors (KIRs) [38].

Regarding the HP group, initially, the high protein intake did not seem to impact pregnancy as significantly as the other diets. However, there was a significant increase in BDA^low^uNK cells with a decrease of pups’ weight after birth. It has been shown that high dietary protein intake has detrimental effects in embryonic development and can lead to low-birth-weight offspring [39]. Nevertheless, there is no data in the literature showing the impact of HP diet on angiogenic uNK cells.

Studies in rats and humans have shown that consumption of HP diets induce hepatic gluconeogenesis [40–42]. During this process, the carbon skeleton of proteins is transformed into carbohydrates or fats to maintain plasma glucose levels [42]. In fact, data from the literature describe the lower anti-inflammatory effects of HP diets due to the amount of saturated fatty acids, which promote inflammation [43]. The increase of inflammatory markers due to consumption of HP diets has also been described [44]. It is plausible that even the HP diet might have induced an increase in DBA^low^uNK cells due to inflammatory mechanisms, albeit on a lesser extent.

In summary, this prospective and translational study sheds light on the importance of uNK cells following a dietary inflammatory process. Further research is essential to elucidate the intricate cellular signaling pathways responsible for the observed changes in DBA^+^uNK cells following dietary consumption. This endeavor holds promise for a comprehensive understanding of the molecular mechanisms underlying the polarization of angiogenic uNK cells in response to nutritional imbalances. Such insights could not only deepen our understanding of reproductive physiology but also pave the way for novel therapeutic interventions targeting uNK cells to mitigate adverse pregnancy outcomes associated with dietary factors.

## Acknowledgments

We thank Dr. Luciano Bruno Carvalho Silva De Carvalho, L. B. for provide all feed used in this study and for valuable suggestions and Dr. Alexandre Giusti-Paiva for assistance in the statistical plan/analysis and valuable suggestions. Funding: This study was financed in part by the Coordenação de Aperfeiçoamento de Pessoal de Nível Superior - Brasil (CAPES) (Finance Code 001) and Fundação de Amparo à Pesquisa de Minas Gerais - FAPEMIG (Proc. No: CBB-APQ-02035-12 and CBB-APQ-01676-13). Salles, E. S. L received a PhD scholarship from CAPES. Da Silva, L. P. received a Ms scholarship from CAPES. DE ALMEIDA, J. G. received an undergraduate scholarship from PIBIC/CNPq.

---

**Table 1.** Centesimal composition of the experimental diets. 1. Values correspond to means (± SD) of three determinations; 2. Values expressed in dry basis; 3. Values not sharing similar letter in the same column are different (p < 0.05) in Tukey test; 4. Calculated by difference = 100 – (protein + total fat + ash + moisture). Control Diet (CD). High Protein Diet (HP). High Fat Diet (HF). High carbohydrate diet (HC) and Food restriction diet (FR).

| Diets | Protein (%)<br>1.2.3 | Fat (%)<br>1.2.3 | Ash (%)<br>1.2.3 | Moisture (%)<br>1.2.3 | Carbohydrate<br>(%)<sup>2,4</sup> | Food<br>availability |
| --- | --- | --- | --- | --- | --- | --- |
| CD | 15.30 ± 0.05<br>c | 7.09 ± 0.06<sup>b</sup> | 2.15 ± 0.08<sup>a</sup> | 7.25 ± 0.05<br>a | 67.21 | *Ad libitum* |
| HP | 31.26 ± 0.17<br>a | 7.07 ± 0.05<sup>b</sup> | 2.18 ± 0.09<sup>a</sup> | 7.15 ± 0.06<br>a | 52.34 | *Ad libitum* |
| HF | 16.55 ± 0.03<br>b | 32.51 ± 0.03<sup>a</sup> | 2.19 ± 0.04<sup>a</sup> | 7.16 ± 0.06<br>a | 41.59 | *Ad libitum* |
| HC | 10.50 ± 0.05<br>d | 2.45 ± 0.03<sup>c</sup> | 2.14 ± 0.09<sup>a</sup> | 7.19 ± 0.03<br>a | 77.72 | *Ad libitum* |
| FR | 15.30 ± 0.05<br>c | 7.09 ± 0.06<sup>b</sup> | 2.15 ± 0.08<sup>a</sup> | 7.25 ± 0.05<br>a | 67.21 | 4g/day |

**Table 1.** Centesimal composition of the experimental diets.
| **Diets** | **Protein (%)**<br>1.2.3 | **Fat (%)**<br>1.2.3 | **Ash (%)**<br>1.2.3 | **Moisture (%)**<br>1.2.3 | **Carbohydrate (%)**<sup>2,4</sup> | **Food availability** |
| --- | --- | --- | --- | --- | --- | --- |
| **CD** | 15.30 ± 0.05<br>c | 7.09 ± 0.06<sup>b</sup> | 2.15 ± 0.08<sup>a</sup> | 7.25 ± 0.05<sup>a</sup> | 67.21 | *Ad libitum* |
| **HP** | 31.26 ± 0.17<br>a | 7.07 ± 0.05<sup>b</sup> | 2.18 ± 0.09<sup>a</sup> | 7.15 ± 0.06<sup>a</sup> | 52.34 | Ad libitum |
| **HF** | 16.55 ± 0.03<br>b | 32.51 ± 0.03<sup>a</sup> | 2.19 ± 0.04<sup>a</sup> | 7.16 ± 0.06<sup>a</sup> | 41.59 | Ad libitum |
| **HC** | 10.50 ± 0.05<br>d | 2.45 ± 0.03<sup>c</sup> | 2.14 ± 0.09<sup>a</sup> | 7.19 ± 0.03<sup>a</sup> | 77.72 | Ad libitum |
| **FR** | 15.30 ± 0.05<br>c | 7.09 ± 0.06<sup>b</sup> | 2.15 ± 0.08<sup>a</sup> | 7.25 ± 0.05<sup>a</sup> | 67.21 | 4g/day |

## Notes

### Competing Interest Statement

The authors have declared no competing interest.

## References

1. Romieu I, Dossus L, Barquera S, Blottiere HM, Franks PW, Gunter M, Hwalla N, Hursting SD, Leitzmann M, Margetts B, Nishida C, Potischman N, Seidell J, Stepien M, Wang Y, Westerterp K, Winichagoon P, Wiseman M, Willett WC, Balance IwgoE, Obesity. Energy balance and obesity: what are the main drivers? Cancer Causes Control 2017; 28: 247–258.

2. Ortigues I, Durand D. Adaptation of energy metabolism to undernutrition in ewes. Contribution of portal-drained viscera, liver and hindquarters. Br J Nutr 1995; 73: 209–26.

3. Kumar PU, Ramalaxmi BA, Venkiah K, Sesikeran B. Effect of maternal undernutrition on human foetal pancreas morphology in second trimester of pregnancy. Indian J Med Res 2013; 137: 302–7.

4. Parker VJ, Solano ME, Arck PC, Douglas AJ. Diet-induced obesity may affect the uterine immune environment in early-mid pregnancy, reducing NK-cell activity and potentially compromising uterine vascularization. Int J Obes (Lond) 2014; 38: 766–74.

5. Ahmed ABA. Association between maternal undernutrition among Sudanese women and newborn birth weight. J Family Med Prim Care 2022; 11: 2824–2827.

6. Bilal JA, Rayis DA, AlEed A, Al-Nafeesah A, Adam I. Maternal Undernutrition and Low Birth Weight in a Tertiary Hospital in Sudan: A Cross-Sectional Study. Front Pediatr 2022; 10: 927518.

7. Deniz A, Okuyucu M. The impact of obesity on fertility and sexual function in women of child bearing age. J Obstet Gynaecol 2022; 42: 3129–3133.

8. Heerwagen MJ, Miller MR, Barbour LA, Friedman JE. Maternal obesity and fetal metabolic programming: a fertile epigenetic soil. Am J Physiol Regul Integr Comp Physiol 2010; 299: R711–22.

9. Ovesen P, Rasmussen S, Kesmodel U. Effect of prepregnancy maternal overweight and obesity on pregnancy outcome. Obstet Gynecol 2011; 118: 305–312.

10. Tabacu MC, Istrate-Ofiteru AM, Manolea MM, Dijmarescu AL, Rotaru LT, Boldeanu MV, Serbanescu MS, Tudor A, Novac MB. Maternal obesity and placental pathology in correlation with adverse pregnancy outcome. Rom J Morphol Embryol 2022; 63: 99–104.

11. Ashkar AA, Croy BA. Functions of uterine natural killer cells are mediated by interferon gamma production during murine pregnancy. Semin Immunol 2001; 13: 235–41.

12. Croy BA, He H, Esadeg S, Wei Q, McCartney D, Zhang J, Borzychowski A, Ashkar AA, Black GP, Evans SS, Chantakru S, van den Heuvel M, Paffaro VA, Jr., Yamada AT. Uterine natural killer cells: insights into their cellular and molecular biology from mouse modelling. Reproduction 2003; 126: 149–60.

13. Peel S. Granulated metrial gland cells. Adv Anat Embryol Cell Biol 1989; 115: 1–112.

14. Delgado SR, McBey BA, Yamashiro S, Fujita J, Kiso Y, Croy BA. Accounting for the peripartum loss of granulated metrial gland cells, a natural killer cell population, from the pregnant mouse uterus. J Leukoc Biol 1996; 59: 262–9.

15. Paffaro VA, Jr., Bizinotto MC, Joazeiro PP, Yamada AT. Subset classification of mouse uterine natural killer cells by DBA lectin reactivity. Placenta 2003; 24: 479–88.

16. Chen Z, Zhang J, Hatta K, Lima PD, Yadi H, Colucci F, Yamada AT, Croy BA. DBA-lectin reactivity defines mouse uterine natural killer cell subsets with biased gene expression. Biol Reprod 2012; 87: 81.

17. Zheng LM, Ojcius DM, Liu CC, Kramer MD, Simon MM, Parr EL, Young JD. Immunogold labeling of perforin and serine esterases in granulated metrial gland cells. FASEB J 1991; 5: 79–85.

18. Croy BA, Reed N, Malashenko BA, Kim K, Kwon BS. Demonstration of YAC target cell lysis by murine granulated metrial gland cells. Cell Immunol 1991; 133: 116–26.

19. Li G, Huang W, Xia Q, Yang K, Liu R, Zhu H, Jiang W. Role of uterine natural killer cells in angiogenesis of human decidua of the first-trimester pregnancy. Sci China C Life Sci 2008; 51: 111–9.

20. Zhang JH, Yamada AT, Croy BA. DBA-lectin reactivity defines natural killer cells that have homed to mouse decidua. Placenta 2009; 30: 968–73.

21. Zavan B, Giusti-Paiva A, Soncini R, do Amarante-Paffaro AM, Paffaro VA, Jr. Immunohistochemical demonstration of blood vessels alpha-actin down-regulation in LPS-treated pregnant mice. Physiol Res 2012; 61: 551–3.

22. Schulz LC, Schlitt JM, Caesar G, Pennington KA. Leptin and the placental response to maternal food restriction during early pregnancy in mice. Biol Reprod 2012; 87: 120.

23. Berg BN. Dietary restriction and reproduction in the rat. J Nutr 1965; 87: 344–8.

24. Pascalon A, Bertrand M. [Effects of overall food restriction on embryo-fetal development in the rat]. Ann Rech Vet 1987; 18: 379–88.

25. Rezaei Kahmini F, Shahgaldi S, Moazzeni SM. Mesenchymal stem cells alter the frequency and cytokine profile of natural killer cells in abortion-prone mice. J Cell Physiol 2020; 235: 7214–7223.

26. Mhd Omar NA, Frank J, Kruger J, Dal Bello F, Medana C, Collino M, Zamaratskaia G, Michaelsson K, Wolk A, Landberg R. Effects of High Intakes of Fructose and Galactose, with or without Added Fructooligosaccharides, on Metabolic Factors, Inflammation, and Gut Integrity in a Rat Model. Mol Nutr Food Res 2021; 65: e2001133.

27. Zavan B, do Amarante-Paffaro AM, Paffaro VA, Jr. alpha-actin down regulation and perforin loss in uterine natural killer cells from LPS-treated pregnant mice. Physiol Res 2015; 64: 427–32.

28. Lv H, Liu Q, Wen Z, Feng H, Deng X, Ci X. Xanthohumol ameliorates lipopolysaccharide (LPS)-induced acute lung injury via induction of AMPK/GSK3beta-Nrf2 signal axis. Redox Biol 2017; 12: 311–324.

29. Wang X, Yang J, Lu T, Zhan Z, Wei W, Lyu X, Jiang Y, Xue X. The effect of swimming exercise and diet on the hypothalamic inflammation of ApoE-/- mice based on SIRT1-NF-kappaB-GnRH expression. Aging (Albany NY) 2020; 12: 11085–11099.

30. Snodgrass RG, Huang S, Choi IW, Rutledge JC, Hwang DH. Inflammasome-mediated secretion of IL-1beta in human monocytes through TLR2 activation; modulation by dietary fatty acids. J Immunol 2013; 191: 4337–47.

31. Yan L, Li Y, Tan T, Qi J, Fang J, Guo H, Ren Z, Gou L, Geng Y, Cui H, Shen L, Yu S, Wang Z, Zuo Z. RAGE-TLR4 Crosstalk Is the Key Mechanism by Which High Glucose Enhances the Lipopolysaccharide-Induced Inflammatory Response in Primary Bovine Alveolar Macrophages. Int J Mol Sci 2023; 24.

32. Chen G, Li J, Ochani M, Rendon-Mitchell B, Qiang X, Susarla S, Ulloa L, Yang H, Fan S, Goyert SM, Wang P, Tracey KJ, Sama AE, Wang H. Bacterial endotoxin stimulates macrophages to release HMGB1 partly through CD14- and TNF-dependent mechanisms. J Leukoc Biol 2004; 76: 994–1001.

33. Cole BK, Morris MA, Grzesik WJ, Leone KA, Nadler JL. Adipose tissue-specific deletion of 12/15-lipoxygenase protects mice from the consequences of a high-fat diet. Mediators Inflamm 2012; 2012: 851798.

34. Herup-Wheeler T, Shi M, Harvey ME, Talwar C, Kommagani R, MacLean JA, 2nd, Hayashi K. High-fat diets promote peritoneal inflammation and augment endometriosis-associated abdominal hyperalgesia. bioRxiv 2023.

35. Orlova EG, Shirshev SV. Leptin as an immunocorrecting agent during normal pregnancy. Bull Exp Biol Med 2009; 148: 75–8.

36. Henson MC, Castracane VD. Leptin in pregnancy. Biol Reprod 2000; 63: 1219–28.

37. Perez-Perez A, Sanchez-Jimenez F, Vilarino-Garcia T, Sanchez-Margalet V. Role of Leptin in Inflammation and Vice Versa. Int J Mol Sci 2020; 21.

38. Castellana B, Perdu S, Kim Y, Chan K, Atif J, Marziali M, Beristain AG. Maternal obesity alters uterine NK activity through a functional KIR2DL1/S1 imbalance. Immunol Cell Biol 2018; 96: 805–819.

39. Whitaker KW, Totoki K, Reyes TM. Metabolic adaptations to early life protein restriction differ by offspring sex and post-weaning diet in the mouse. Nutr Metab Cardiovasc Dis 2012; 22: 1067–74.

40. Azzout-Marniche D, Gaudichon C, Blouet C, Bos C, Mathe V, Huneau JF, Tome D. Liver glyconeogenesis: a pathway to cope with postprandial amino acid excess in high-protein fed rats? Am J Physiol Regul Integr Comp Physiol 2007; 292: R1400–7.

41. Veldhorst MA, Westerterp-Plantenga MS, Westerterp KR. Gluconeogenesis and energy expenditure after a high-protein, carbohydrate-free diet. Am J Clin Nutr 2009; 90: 519–26.

42. Pesta DH, Samuel VT. A high-protein diet for reducing body fat: mechanisms and possible caveats. Nutr Metab (Lond) 2014; 11: 53.

43. Koelman L, Markova M, Seebeck N, Hornemann S, Rosenthal A, Lange V, Pivovarova-Ramich O, Aleksandrova K. Effects of High and Low Protein Diets on Inflammatory Profiles in People with Morbid Obesity: A 3-Week Intervention Study. Nutrients 2020; 12.

44. Hruby A, Jacques PF. Dietary Protein and Changes in Biomarkers of Inflammation and Oxidative Stress in the Framingham Heart Study Offspring Cohort. Curr Dev Nutr 2019; 3: nzz019.

