## Supplementary figure 1 for "The role of dietary patterns in the polarization of angiogenic uterine Natural Killer cells during murine pregnancy"

| Composition (%) | CD | HP | HL | HC |
|---|---|---|---|---|
| Casein | 20,00 | 42,00 | 20,00 | 15,00 |
| L-Cystine | 0,30 | 0,30 | 0,30 | 0,30 |
| Corn starch | 39,75 | 17,75 | 13,75 | 49,75 |
| α-Corn starch | 13,20 | 13,20 | 13,20 | 13,20 |
| Sucrose | 10,00 | 10,00 | 10,00 | 10,00 |
| Soybean oil | 7,00 | 7,00 | 4,00 | 2,00 |
| Cellulose powder | 5,00 | 5,00 | 5,00 | 5,00 |
| AIN-93G mineral mix | 3,50 | 3,50 | 3,50 | 3,50 |
| AIN-93G vitamin mix | 1,00 | 1,00 | 1,00 | 1,00 |
| Heavy tartaric acid choline | 0,25 | 0,25 | 0,25 | 0,25 |
| Tertiary butyl hydroquinone | 0,0014 | 0,0014 | 0,0014 | 0,0014 |
| Hydrogenated vegetable oil (unsaturated fat) | 0 | 0 | 29,00 | 0 |
| **Total** | 100 | 100 | 100 | 100 |

**Supplementary figure 1**. Detailed composition of control (CD), High protein (HP), High Fat (HF), High carbohydrate (HC) foods (g/100g).

**a**

| | SI | SII | SIII | SIV | DBA$^{low}$ |
|---|---|---|---|---|---|
| **CD** | 15.2 ± 7.7 | 15.7 ± 5 | 2.9 ± 3.2 | 0.02 ± 0.05 | 0.13 ± 0.18 |
| **HP** | 10.9 ± 7.8 | 11.8 ± 3.6 | 1.6 ± 1.7 | 3.8 ± 4.3* | 5.3 ± 2.8 |
| **HF** | 13.2 ± 6.5 | 5.2 ± 4.5** | 0.1 ± 0.16* | 0.2 ± 0.3 | 15.6 ± 9.5*** |
| **HC** | 9 ± 4.7 | 1.6 ± 1.6*** | 0.02 ± 0.05* | 0.0 ± 0.0 | 27.4 ± 10*** |
| **FR** | 17.1 ± 8 | 14.3 ± 6.1 | 0.1 ± 0.14* | 0.04 ± 0.1 | 0 ± 0 |

**b**

| | SI | SII | SIII | SIV | DBA$^{low}$ |
|---|---|---|---|---|---|
| **CD** | 1.02 ± 1.5 | 4.04 ± 2.4 | 14.2 ± 3.5 | 2.3 ± 1.3 | 0.2 ± 0.2 |
| **HP** | 0.16 ± 0.15 | 2.18 ± 1.15 | 3.8 ± 0.5*** | 9.6 ± 3.2* | 7 ± 2.1* |
| **HF** | 0.09 ± 0.09 | 0.64 ± 0.7 | 0.95 ± 1.0*** | 1.7 ± 0.9 | 21.4 ± 7.8**** |
| **HC** | 0.07 ± 0.10 | 0.6 ± 0.5 | 0.78 ± 0.74*** | 1.2 ± 1 | 16.4 ± 3.4*** |
| **FR** | 1,97 ± 2.14 | 13.5 ± 5.9*** | 13 ± 6.4 | 1.7 ± 1 | 0.3 ± 0.4 |

**c**

| | SI | SII | SIII | SIV | DBA$^{low}$ |
|---|---|---|---|---|---|
| **CD** | 0.09 ± 0.09 | 1.16 ± 1 | 9.4 ± 3.4 | 11.5 ± 4.3 | 0.18 ± 0.19 |
| **HP** | 0.04 ± 0.06 | 1.1 ± 0.8 | 2 ± 1*** | 17 ± 4.3*** | 6.2 ± 2.2*** |
| **HF** | 0.04 ± 1 | 0.2 ± 0.1 | 0.7 ± 0.7*** | 6.9 ± 4.8 | 21.9 ± 3.1*** |
| **HC** | 0.04 ± 0.06 | 0.04 ± 0.06 | 0.15 ± 0.2*** | 7.8 ± 2.7 | 14 ± 4.4*** |
| **FR** | 1 ± 1.4 | 7.9 ± 4.4*** | 16.3 ± 5.7*** | 4.5 ± 3* | 0.4 ± 0.5 |

**d**

| | SI | SII | SIII | SIV | DBA$^{low}$ |
|---|---|---|---|---|---|
| **CD** | 8.1 ± 1.1 | 12.8 ± 0.8 | 23.5 ± 1 | 27.5 ± 1 | 23.1 ± 0.7 |
| **HP** | 10 ± 0.5** | 13.8 ± 1 | 23.4 ± 1 | 26.5 ± 1.9 | 21.8 ± 2 |
| **HF** | 9.2 ± 0.5 | 13.7 ± 1 | 24.3 ± 1.9 | 24.6 ± 2 | 25.3 ± 1.4 |
| **HC** | 9.9 ± 0.6* | 14 ± 1.7 | 25 ± 1.8 | 25 ± 2.5 | 24.4 ± 1.8 |
| **FR** | 8.5 ± 1 | 11 ± 2 | 18.3 ± 2.9** | 19.4 ± 2.6*** | 17.3 ± 0.8*** |

**Supplementary figure 2**. Uterine Natural Killer cells stereological and Morphometrical analyses. Subtypes quantification at Region 1 (A). Subtypes quantification at Region 2 (B). Subtypes quantification at Region 3 (C). Subtype uNK diameter (D).
